# A single nucleotide polymorphism of *PLIN2* is associated with nonalcoholic steatohepatitis and causes phenotypic changes in hepatocyte lipid droplets

**DOI:** 10.1101/823450

**Authors:** Claire S Faulkner, Collin White, Loretta L Jophlin

**Affiliations:** University of Nebraska Medical Center Department of Internal Medicine, Omaha, NE; Washington University, St. Louis, MO; Mayo Clinic, Division of Gastroenterology and Hepatology, Rochester, MN

**Author notes:** **Corresponding Author:** Loretta L Jophlin 200 1^st^ St SW Rochester, MN 55905. **Conflict of Interest Statement:** The authors have nothing to disclose. **Funding Sources:** LLJ was supported by the UNMC Internal Medicine Development Funds and the UNMC College of Medicine Program of Excellence Physician-Scientist Training Program.

**Keywords:** Non-alcoholic steatohepatitis, perilipin 2, hepatic steatosis, lipid droplet, single nucleotide polymorphism

## Abstract

Perilipin 2 (PLIN2) is a lipid droplet-associated protein which regulates cellular lipid storage and is highly expressed in the liver. Previous studies of a missense single nucleotide polymorphism (SNP) in *PLIN2*, Ser251Pro, (i.e. rs3556875) have shown this SNP to cause decreased lipolysis and increased intracellular lipid accumulation. To explore if this SNP is associated with nonalcoholic steatohepatitis (NASH), we genotyped 116 adults with NASH and 67 age- and gender-matched controls. rs3556875 was significantly associated with NASH with an allelic odds ratio of 2.98 (95% confidence interval 1.12 - 7.31, *p*=0.02) and in a dominant inheritance model. In an *in vitro* model of hepatic steatosis, expression of the Pro251 variant protein led to phenotypic changes in intracellular lipid droplets, yielding smaller diameter and up to 7 fold more lipid droplets per cell when compared to wild type *PLIN2*. In conclusion, the Ser251Pro SNP of *PLIN2* may convey risk for NASH and causes phenotypic changes in hepatocyte lipid droplets.

## INTRODUCTION

Nonalcoholic steatohepatitis (NASH) cirrhosis is a leading etiology of liver disease worldwide, paralleling the rise in the metabolic syndrome (1). While non-alcoholic fatty liver disease affects approximately 25% of the global population (2), progression to NASH occurs less frequently and varies amongst individuals due to comorbid conditions and genetic risk factors (3). Two recent population studies (4, 5) implicate a single nucleotide polymorphism (SNP) in the perilipin 2 (*PLIN2*) gene, rs35568725, in atherosclerosis and insulin resistance. This SNP codes a Ser251Pro variant and is associated with smaller, more numerous lipid droplets (LD) and reduced lipolysis in macrophages (4). Germline and hepatic loss of *PLIN2* have been shown to abrogate hepatic steatosis in animal models of NASH (6) but investigations of the *PLIN2* Ser251Pro variant have not been reported in the context of liver disease.

Our study is the first to investigate rs35568725 in NASH, a disease driven by altered hepatocyte LD metabolism. We determined the prevalence of rs35568725 in a population of adults with and without NASH. To determine if the Pro251 variant of *PLIN2* affects LD phenotype, we assessed hepatocyte LDs containing the wild type and Pro251 variants within an *in vitro* model of hepatic steatogenesis. Our findings suggest rs35568725 conveys risk for NASH in humans and that the mechanism may be due to increased hepatocyte LD burden under steatogenic conditions.

## MATERIALS AND METHODS

The Nebraska Biobank, a biorepository with human genomic DNA and linked deidentified health data was utilized. Subjects gave written informed consent and the study was approved by the University of Nebraska Medical Center Institutional Review Board. All subjects age ≥ 18 with a diagnosis of NASH excluding other etiologies of liver disease (viral, alcohol, autoimmune, etc.) were included (n=116). Using previous population studies of rs35568725 with minor allele frequencies of 0.06, 0.05 (4), and 0.03 (5), and a 5% margin of error with a 95% confidence interval, a sample size of 67 age- and gender-matched controls lacking NASH and other liver disease was deemed appropriate for this pilot study.

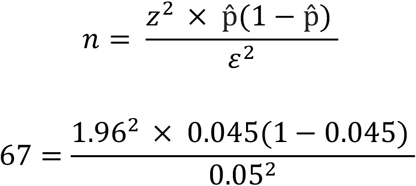

Genomic DNA was extracted from blood with the DNA QIAcube HT kit (Qiagen, Germantown, MD USA). PCR of the *PLIN2* region of interest was performed using forward (5’AGGTTAGAGTCCAGGCCTTAT3’) and reverse (5’GAATATGGAGACAGCTCACAGAA3’) primers, yielding a 408 bp product verified by agarose gel. Amplicons were purified using ExoSAP-IT™ PCR Cleanup Reagent (ThermoFisher Scientific, Waltham, MA USA) and sequenced using Applied Biosystems 3730 DNA Analysis Instrument (Life Technologies, Grand Island, NY USA). Two authors (CSF and LLJ) remained blinded to subject status and used SnapGene (GSL Biotech, Chicago, IL USA) to determine genotypes.

### *In vitro* phenotype of PLIN2 variants

Using site-directed mutagenesis (QuikChange Lightning, Agilent, Santa Clara, CA USA), the Pro251 allele was inserted into the green fluorescent protein (GFP)-PLIN2 plasmid (Addgene #87161). Sequences were verified in forward and reverse directions using primers EGFP-N and SV40pA-R, respectively. Huh7 hepatocytes were reverse transfected with efficiencies > 50% using Lipofectamine2000 (ThermoFisher Scientific, Waltham, MA USA) and steatosis was induced for 48 hours with 0.8mM oleic acid as described (7). Comparative morphometrics of LDs was performed on three independent experiments using confocal (Zeiss LSM 800 with Airyscan, Carl Zeiss, Obercohen, Germany) optical microscopy (Echo Revolve, San Diego, CA USA) and ImageJ software as described (8). LD measurements are expressed as mean +/− one standard deviation.

### Genetic models and statistical analysis

The Hardy–Weinberg equilibrium, allelic odds ratio (OR) and genotypic OR using dominant and recessive models were calculated as reviewed (9). To determine differences between groups for all variables, T-test, Mann-Whitney U and Chi-square test were used as indicated. P values <0.05 were deemed significant. All analyses were performed using GraphPadPrism 8.1.2 (GraphPad Software, La Jolla, CA USA).

## RESULTS

Control and NASH groups were matched for gender and age and discordant for comorbidities associated with NASH as expected (supplemental table 1). rs35568725 was in Hardy–Weinberg equilibrium (χ^2^ 3.52, p<0.05) in our subject population. Of 116 NASH subjects, 81.9% (n=95) lacked, 15.5% (n=18) had a single copy, and 2.6% (n=3) had two copies of rs35568725. Of 67 controls, 92.5% (n=62) lacked, 7.5% (n=5) had one copy and none were homozygous for rs35568725. Due to its lower frequency, rs35568725 was deemed to be the minor allele (a) and wild type was deemed the major allele (A). Allelic OR for NASH conveyed by the minor allele was found to be 2.98 (1.12 – 7.31, p=0.02) (Table 1). In a dominant inheritance model, the OR for NASH conveyed by the minor allele was 2.74 (1.01-6.92, p=0.04) (Table 1). Within the NASH cohort, rs35568725 carriers did not have higher rates of NASH risk factors including obesity (body mass index >30), type 2 diabetes and hyperlipidemia or cirrhosis compared to non-carriers (supplemental Table 2).

**Table 1.** Genetic models of association between rs35568725 and NASH

Genetic models of association between rs35568725 and NASH
| Genetic Model | OR (95% CI) | p value<sup>1</sup> |
| --- | --- | --- |
| ***Allelic: (a vs. A)*** | 2.98 (1.12 – 7.31) | **0.02** |
| **Dominant: (Aa + aa) vs. AA** | 2.74 (1.01 – 6.92) | **0.04** |
| **Recessive: (aa) vs. Aa + AA** | Indeterminate | Indeterminate |

<sup>1</sup>Chi-square test, A: Wild type, a: rs35568725, OR: odds ratio, CI: confidence interval

Within transfected Huh7 hepatocytes, overexpressed GFP-PLIN2 and GFP-PLIN2Pro251 localized to LDs. Under basal conditions, hepatocytes expressing GFP-PLIN2Pro251 compared to GFP-PLIN2 showed similar diameter LDs (0.59 +/− 0.35 vs 0.71 +/− 0.51µm, *p*=0.09) and number of LDs per cell (93 +/− 41.68 vs 45.33 +/− 10.69 *p*=0.13) (Figure 1). Under steatogenic conditions however, hepatocytes expressing GFP-PLIN2Pro251 had smaller diameter (0.89 +/− 0.27 vs 1.49 +/− 0.76 µm, *p*=<0.0001) and more numerous LDs per cell (362.7 +/− 131.2 vs 53.67 +/− 40.15, *p*=0.0175) than did GFP-PLIN2 expressing hepatocytes (Figure 1).

**Figure 1.**
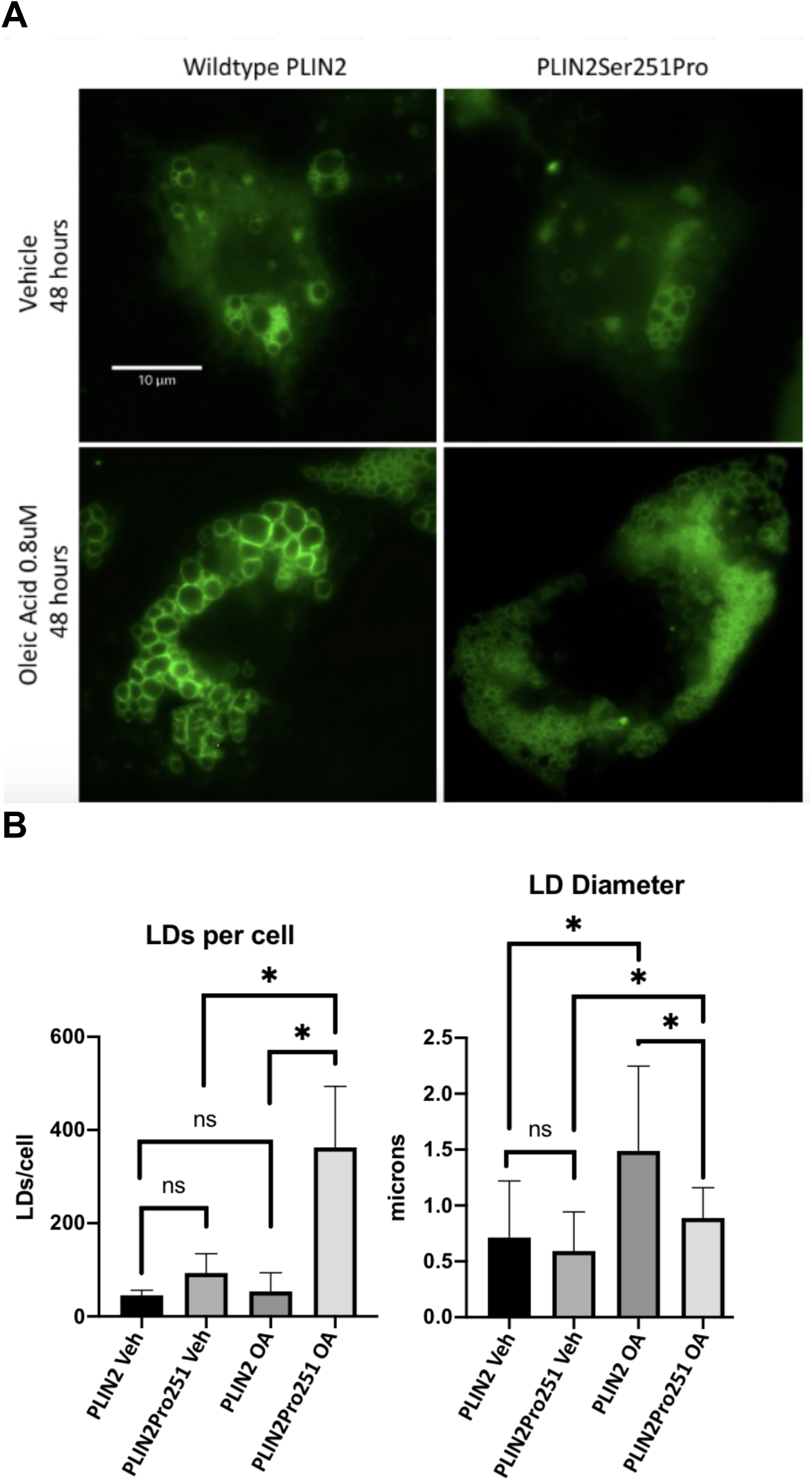
(A) Fluorescence microscopy of representative single Huh7 hepatocytes expressing green fluorescent protein tagged (GFP)-PLIN2 (left) and GFP-PLIN2Pro251 (right) localized to lipid droplets (LD) treated with vehicle (top) and 0.8 mM oleic acid for 48 hours (bottom). (B) LDs per cell (left) and LD diameter (right) quantified from ≥10 cells from 3 independent experiments (N=3). Veh = vehicle treatment, OA = oleic acid 0.08 mM, Scale bar = 10 um, Error Bar = one standard deviation, * p<0.05, ns not significant.

## DISCUSSION

Within a discovery cohort, we found rs35568725 may convey risk for NASH. In our population using a dominant inheritance model, subjects with rs35568725 were 2.74-fold more likely to have NASH. This SNP may be an independent risk factor for NASH because within the NASH group, subjects with rs35568725 had similar frequencies of typical NASH comorbidities as subjects without the SNP. rs35568725 may not portend a more aggressive trajectory of liver disease, as carriers had the same rate of cirrhosis as non-carriers in the NASH group. Limitations of this pilot study are the racial homogeneity of our population and relatively small sample size. No subjects in our control cohort were homozygous for rs35568725 precluding use of additional genotypic models of inheritance (e.g. recessive).

Our *in vitro* work herein reveals the impact that the PLIN2Pro251 variant protein has on hepatocyte LD phenotype. Under basal conditions, hepatocyte LDs harboring wild type PLIN2 and PLIN2Pro251 are similar in size and number. Under steatogenic conditions however, hepatocytes expressing PLIN2Pro251 become heavily laden with small LDs, accumulating up to ~7 fold more LDs that are approximately half the diameter of LDs circumscribed with wild type PLIN2. LDs in PLIN2Pro251 expressing hepatocytes also appear more homogeneous in phenotype, with less size variability than those in wild type PLIN2 expressing hepatocytes. As PLIN2 plays regulatory roles in many aspects of LD biology, we are undertaking a comprehensive mechanistic assessment of PLIN2Pro251. Others have shown that PLIN2Pro251 decreases lipolysis in macrophages (4), and we are investigating the enzymatic step of lipolysis impacted by the variant protein. While the 3-dimensional structure of PLIN2 is not known, based on homology with the structure of perilipin 3 (10), it is theorized that the Pro251 substitution causes a tertiary change in PLIN2, which may decrease hormone sensitive lipase (11) or adipose triglyceride lipase (12) docking to PLIN2 on the LD surface. Further research is needed to delineate if the Pro251 variant impacts aspects of LD biology outside of lipolysis including biogenesis in the endoplasmic reticulum (13), fusion (14) and autophagy (12) summarized in figure 2.

**Figure 2.**
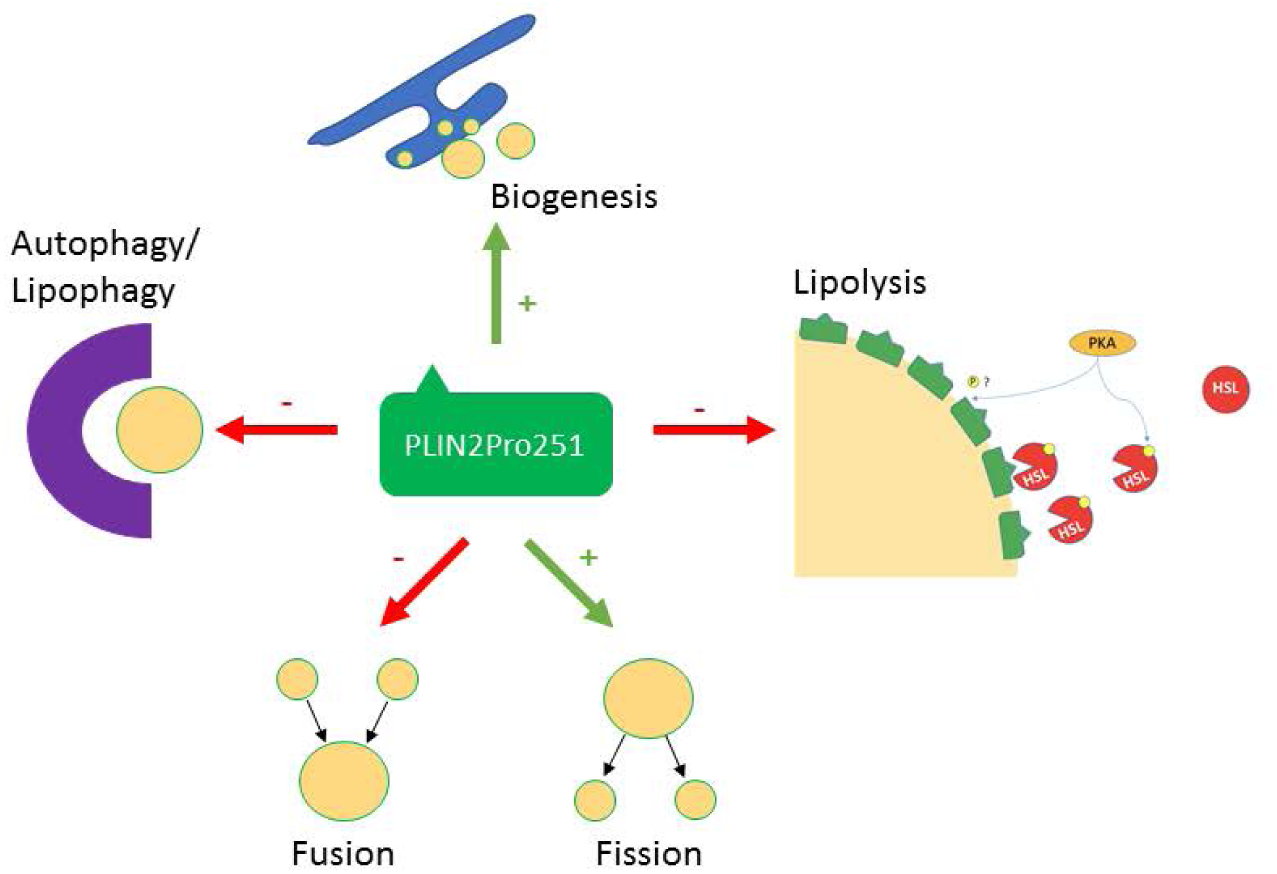
Possible mechanisms for PLIN2Pro251 phenotypic influence on lipid droplets. PKA = protein kinase A, HSL = hormone sensitive lipase.

In summary, we demonstrate that a coding SNP of *PLIN2*, rs35568725, may convey risk for NASH in a discovery cohort. When expressed under steatogenic conditions, human hepatocytes harboring the PLIN2Pro251 variant protein become heavily laden with small, homogenous LDs. More work is ongoing to validate rs35568725 as an independent risk factor for NASH in broader, more diverse populations and to delineate the mechanism(s) of its effects on LD phenotype.

## Acknowledgements

We thank Mary Anne Phillippi and Cody Wehrkamp (University of Nebraska Medical Center) for assistance with PCR and cell culture. We acknowledge U54GM115458/NIGMS NIH HHS/United States, the Great Plains IDeA-CTR grant and the Nebraska Research Initiative, which subsidizes the Nebraska Biobank repository, the UNMC Advanced Microscopy Core Facility and Center for Cellular Signaling CoBRE “NIH P30GM106397” *a*nd the UNMC DNA Sequencing Core Facility, which receives partial support from the Nebraska Research Network In Functional Genomics NE-INBRE P20GM103427-14, the Molecular Biology of Neurosensory Systems CoBRE P30GM110768, the Fred & Pamela Buffett Cancer Center – P30CA036727, the Center for Root and Rhizobiome Innovation (CRRI) 36-5150-2085-20, and the Nebraska Research Initiative.

---

**Supplemental table 1.** Clinical characteristics of control and NASH subjects

**All subjects**
|  | **Control**<br>**(n=67)** | **NASH**<br>**(n=116)** | **p value<sup>1</sup>** |
| --- | --- | --- | --- |
| *Age (Years)* | 59.71 | 56.02 | 0.32 |
| *Gender (Female)* | 34 (50.75%) | 68 (58.62%) | 0.35<sup>‡</sup> |
| *Race (Caucasian)* | 59 (88.06%) | 102 (87.93%) | >0.992<sup>‡</sup> |
| *BMI (kg/m<sup>2</sup>)* | 27.93 | 34.80 | **<0.01** |
| *Hemoglobin A1C (%)* | 5.44 | 7.03 | **<0.01** |
| *Albumin (g/dL)* | 3.96 | 3.71 | **<0.01** |
| *Alanine Aminotransferase (U/L)* | 18.63 | 70.20 | **<0.01** |
| *Alkaline Phosphatase (U/L)* | 73.87 | 105.80 | **<0.01** |
| *Aspartate Aminotransferase (U/L)* | 26.02 | 64.80 | 0.12 |
| *Bilirubin (mg/dL)* | 0.66 | 1.44 | **0.04** |
| *Total Cholesterol (mg/dL)* | 172.30 | 174.20 | 0.85 |
| *Creatinine (mg/dL)* | 0.91 | 1.15 | 0.07 |
| *Ferritin (units)* | 230.60 | 325.80 | 0.57 |
| *Hemoglobin (g/dL)* | 13.50 | 12.90 | 0.12 |
| *LDL (mg/dL)* | 99.35 | 95.96 | 0.67 |
| *Sodium (mmol/L)* | 138.60 | 137.30 | 0.10 |
| *INR* | 1.17 | 1.20 | 0.77 |
| *Platelets (x10<sup>3</sup>/uL)* | 220.20 | 187.80 | **0.03** |
| *Protein (g/dL)* | 6.95 | 7.00 | 0.72 |
| *Triglycerides (mg/dL)* | 115.10 | 181.70 | **0.03** |
| *HDL (mg/dL)* | 49.44 | 38.06 | **<0.01** |

**Control subjects by genotype**
|  | **AA**<br>**(n=62)** | **Aa**<br>**(n=5)** | **p value<sup>2</sup>** |
| --- | --- | --- | --- |
| *BMI (kg/m<sup>2</sup>)* | 28.14 | 25.47 | 0.27 |
| *Hemoglobin A1C (%)* | 5.30 | 6.10 | 0.08 |
| *Albumin (g/dL)* | 4.00 | 4.15 | 0.53 |
| *Alanine Aminotransferase (U/L)* | 18.00 | 18.50 | 0.77 |
| *Alkaline Phosphatase (U/L)* | 74.12 | 71.00 | 0.80 |
| *Aspartate Aminotransferase (U/L)* | 23.00 | 28.00 | **0.04** |
| *Bilirubin (mg/dL)* | 0.53 | 0.55 | 0.90 |
| *INR* | 1.14 | Data not available |  |
| *Platelets (x10<sup>3</sup>/uL)* | 218.00 | 255.00 | 0.21 |
| *Triglycerides (mg/dL)* | 118.80 | Data not available |  |
| *HDL (mg/dL)* | 48.54 | Data not available |  |

**NASH subjects by genotype**
|  | **AA**<br>**(n=95)** | **Aa/aa**<br>**(n=21)** | **p value<sup>1</sup>** |
| --- | --- | --- | --- |
| *BMI (kg/m<sup>2</sup>)* | 34.74 | 35.08 | 0.87 |
| *Hemoglobin A1C (%)* | 7.05 | 6.95 | 0.86 |
| *Albumin (g/dL)* | 3.69 | 3.76 | 0.65 |
| *Alanine Aminotransferase (U/L)* | 54.45 | 140.70 | **0.02** |
| *Alkaline Phosphatase (U/L)* | 102.50 | 120.20 | 0.09 |
| *Aspartate Aminotransferase (U/L)* | 69.96 | 41.67 | 0.52 |
| *Bilirubin (mg/dL)* | 1.46 | 1.32 | 0.83 |
| *INR* | 1.20 | 1.14 | 0.65 |
| *Platelets (x10<sup>3</sup>/uL)* | 186.10 | 195.70 | 0.67 |
| *Triglycerides (mg/dL)* | 187.00 | 155.30 | 0.41 |
| *HDL (mg/dL)* | 37.98 | 39.25 | 0.89 |

A: Wild type, a: rs35568725, <sup>1</sup>T-test, <sup>‡</sup>Chi-square test, <sup>2</sup>Mann-Whitney U

**Supplemental Table 2.** Genotype and comorbidities within the NASH subject cohort

| **Comorbidities** | **AA**<br>**(n=95)** | **Aa or aa**<br>**(n=21)** | **p value<sup>1</sup>** |
| --- | --- | --- | --- |
| *Type 1 Diabetes Mellitus* | 2.11% | 0% | >0.99 |
| *Type 2 Diabetes Mellitus* | 58.95% | 52.38% | 0.63 |
| *Obesity (BMI≥30)* | 67% | 71% | 0.87 |
| *Hyperlipidemia* | 42.70% | 52.40% | 0.48 |
| *Cirrhosis* | 36.84% | 38.10% | >0.99 |
| *Ascites* | 23.16% | 23.81% | >0.99 |
| *Portosystemic Encephalopathy* | 23.16% | 23.81% | >0.99 |
| *Varices* | 14.74% | 23.81% | 0.33 |
| *Hepatocellular Carcinoma* | 3.16% | 0% | >0.99 |
| *Liver Transplant* | 12.63% | 4.76% | 0.46 |

